# TRPV4 channel contributes to aortic root stiffening and atherosclerotic lesion development

**DOI:** 10.1101/2024.11.07.622110

**Authors:** Suneha G. Rahaman, Bidisha Dutta, Shaik O. Rahaman

## Abstract

Cardiovascular disease is the number one cause of death in the developed world and atherosclerosis, a chronic arterial disease, is the most dominant underlying pathology. Epidemiologic and experimental studies suggest that arterial stiffness is a risk factor for atherosclerosis. However, there has been surprisingly limited development in mechanistic understanding of the generation of arterial stiffness and little progress in understanding the mechanisms by which matrix stiffening drives the development of atherosclerosis.

Various proinflammatory and fibrotic activities of macrophages and fibroblasts, such as migration, inflammatory gene expression, and myofibroblast activation, are influenced by matrix stiffness. This influence suggests that aorta stiffening may regulate atherosclerosis via a cellular stiffness sensor. Our research indicates that mechanosensitive transient receptor potential vanilloid 4 (TRPV4) channels control inflammation and fibrosis in other organs and regulate macrophage and fibroblast activation, implicating TRPV4 as a potential stiffness sensor in atherosclerosis. This suggests a cycle where inflammation, fibrosis, and tissue stiffening reinforce each other, with macrophages playing a key role. Here, we identify a cellular stiffness sensor linking matrix stiffness and atherosclerosis using human aortic tissues, a murine atherosclerosis model, and atomic force microscopy (AFM) analysis. This novel finding suggests that targeting TRPV4 could be a selective strategy to prevent or suppress atherogenesis.

## Introduction, Results, and Discussion

Cardiovascular disease is the number one cause of death in the developed world and atherosclerosis, a chronic arterial disease, is the most dominant underlying pathology (1). Epidemiologic and experimental studies suggest that arterial stiffness is a risk factor for atherosclerosis (2, 3). However, there has been surprisingly limited development in mechanistic understanding of the generation of arterial stiffness and little progress in understanding the mechanisms by which matrix stiffening drives the development of atherosclerosis.

Various proinflammatory and fibrotic activities of macrophages and fibroblasts, such as migration, inflammatory gene expression, and myofibroblast activation, are influenced by matrix stiffness. This influence suggests that aorta stiffening may regulate atherosclerosis via a cellular stiffness sensor (4, 5). Our research indicates that mechanosensitive transient receptor potential vanilloid 4 (TRPV4) channels control inflammation and fibrosis in other organs and regulate macrophage and fibroblast activation, implicating TRPV4 as a potential stiffness sensor in atherosclerosis (4, 5). This suggests a cycle where inflammation, fibrosis, and tissue stiffening reinforce each other, with macrophages playing a key role. Here, we identify a cellular stiffness sensor linking matrix stiffness and atherosclerosis using human aortic tissues, a murine atherosclerosis model, and atomic force microscopy (AFM) analysis.

The effect of TRPV4 on atherosclerosis was studied in a well-established *ApoE*^*-/-*^ mouse model of atherosclerosis compared to *ApoE*^*-/-*^*Trpv4*^*-/-*^ mice (both on a C57BL/6 background; n = 10 mice/group) using high-fat diet (HFD) (21% fat, 0.15% cholesterol, no cholate; Harlan Teklad, Madison, WI) and control chow diet. We found that after 12 weeks on a HFD *ApoE*^*-/-*^*Trpv4*^*-/-*^ mice exibited a 4-fold reduction in the amount of atherosclerotic plaque in the aortic root and aortic arch areas compared to *ApoE*^*-/-*^ mice (Figure A). There were no significant differences in body weight gain, plasma total cholesterol (1120 ± 200 vs 1089 ± 170 mg/dL) or triglyceride (120 ± 11 vs 116 ± 12 mg/dL) levels between *ApoE*^*-/-*^*Trpv4*^*-/-*^ and *ApoE*^*-/-*^ mice, suggesting that the observed atheroprotective effect may be linked to the absence of TRPV4-dependent functions in this model. To test TRPV4’s role in regulating aortic root stiffness in vivo, we measured stiffness using AFM on 10-μm sections from *ApoE*^*-/-*^ and *ApoE*^*-/-*^*Trpv4*^*-/-*^ mice. Results showed a 6-fold increase in aortic root stiffness in *ApoE*^*-/-*^ mice on a HFD compared to a chow diet (Figure B). Additionally, *ApoE*^*-/-*^ mice had a 3-fold higher stiffness than *ApoE*^*-/-*^*Trpv4*^*-/-*^ mice on HFD, indicating TRPV4’s involvement in aortic stiffness during atherogenesis (Figure B). Atherogenesis involves aortic stiffening due to extracellular matrix remodeling and cellular changes (2, 3), although the exact mechanism remains unknown.

**Figure 1.**
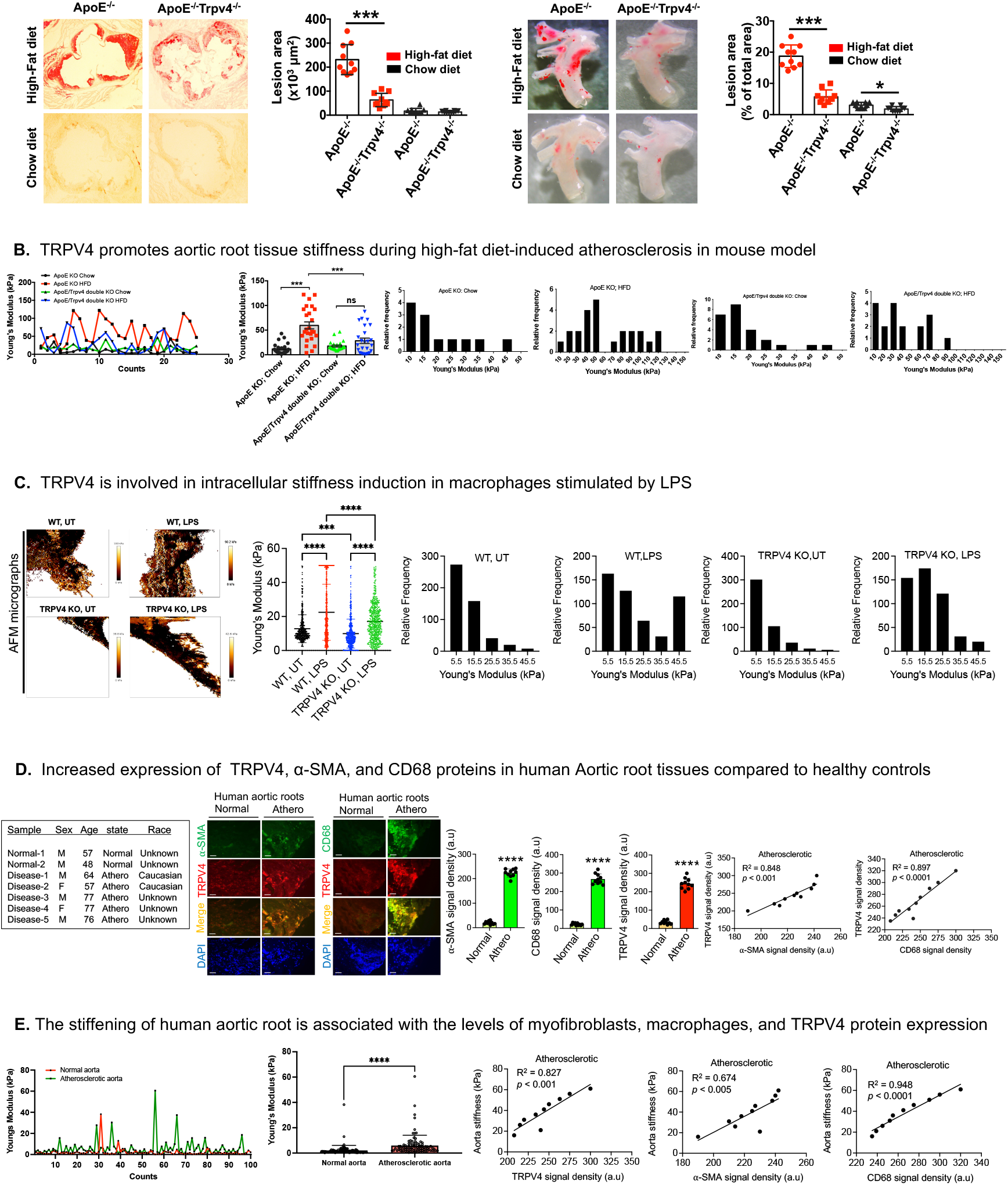
TRPV4 regulates matrix stiffening in atherosclerosis. **A**. Six-week-old male *ApoE*^*-/-*^*Trpv4*^*-/-*^ and control *ApoE*^*-/-*^ mice were maintained on HFD or chow diet for 12 weeks. Lesion areas of Oil-red-O-stained plaque in aortic sinus and arch areas were analyzed. Quantitation of results were shown in bar graphs. n = 10 mice/group, *p ≤ 0.05, ***p ≤ 0.001, t test. **B**. Plot profile of Young’s modulus (YM) of the dataset (*ApoE*^*-/-*^ vs. *ApoE*^*-/-*^*Trpv4*^*-/-*^ mice; HFD or chow for 12 weeks) generated by AFM analysis. n = 5 tissue samples/group and 30 force curves/sample; Quantification of YM of the dataset. n = 5 tissue samples/group and 30 force curves/sample. \*\*\**p* < 0.001, ANOVA; Representative frequency distributions of force curves were shown. **C**. AFM micrographs to represent variations in stiffness in WT and TRPV4 KO macrophages (untreated (UT) or LPS (100 ng/ml) for 24 h); Quantification of YM of the dataset generated by AFM analysis. n = 5 cells/group and 50 force curves/sample. \*\*\**p* < 0.001, ANOVA; Representative frequency distributions of force curves. **D**. Clinical parameters of atherosclerosis and normal subjects; Immunofluorescence staining of aortic root sections from patients or healthy-subjects to identify α-SMA+ myofibroblasts (anti-CD68; 1:100; Sigma-Aldrich) or CD68+ macrophages (anti-CD68; 1:100; Bio-Rad) expressing TRPV4 proteins (anti-TRPV4; 1:100; Alomone, Israel). Quantitation of results show expression of α-SMA, CD68, and TRPV4 (n = 5 sections/condition; \*\*\*\**p* < 0.0001, ANOVA); Scatterplot with linear regression analysis shows TRPV4 signals relative to α-SMA or CD68 abundance in five atherosclerotic root samples. **E**. Plot profile of YM of the dataset (atherosclerotic-root vs. normal-root) generated by AFM analysis. Quantification of YM of the dataset. n = 2-5 tissue samples/group and 50 force curves/sample. \*\*\*\**p* < 0.0001, ANOVA. Scatterplot with linear regression analysis shows the atherosclerotic aortic root stiffness relative to α-SMA, TRPV4, or CD68 abundance in five atherosclerotic samples.

We tested TRPV4’s role in macrophage, an essential cell in atherogenesis, stiffness using AFM and found increased stiffness in WT macrophages compared to TRPV4 KO macrophages. This occurred with both unstimulated and LPS-stimulated, suggesting TRPV4’s involvement in macrophage stiffness during atherogenesis (Figure C).

We presented clinical parameters of atherosclerotic and healthy subjects in Figure D. To detect TRPV4, CD68, and α-smooth muscle actin (α-SMA) proteins in human aortic root tissues, we obtained 5-μm sections of aortic root biopsies from atherosclerosis patients (n = 5) and healthy subjects (n = 2) (OriGene; Rockville, MD). Double immunofluorescence staining identified α-SMA+ myofibroblasts and CD68+ macrophages expressing TRPV4 proteins (TRPV4+). Immunofluorescence intensity was quantified, showing increased α-SMA+ myofibroblasts and CD68+ macrophages in atherosclerotic aortic roots compared to healthy controls, with both cell types coexpressing TRPV4 (Figure D). We observed an ∼8-fold increase in TRPV4, CD68, and α-SMA expression in atherosclerotic aortic roots versus healthy controls (Figure D). Additionally, we found a strong correlation between TRPV4 and CD68 or α-SMA expression, suggesting TRPV4+-myofibroblasts and -macrophages are present in human atherosclerotic aortic tissues (Figure D). This aligns with our hypothesis that TRPV4 drives myofibroblast differentiation and macrophage accumulation in atherosclerosis, supported by previous work showing TRPV4’s role in macrophage accumulation in fibrotic skin tissues (4, 5).

To test whether aortic root tissue stiffening is associated with human atherosclerosis, we measured aortic root stiffness (Young’s modulus) using AFM (5). Five-μm aortic root sections from atherosclerotic patients and controls were analyzed to obtain force curves. We found aortic root tissue stiffness to be ∼4-fold higher in atherosclerotic root compared to healthy roots, indicating increased stiffness in atherosclerotic aortic roots (Figure E). Additionally, analysis showed a strong correlation between root stiffness and levels of myofibroblasts (α-SMA+), macrophages (CD68+), and TRPV4 proteins, suggesting a role for these cells in aortic tissue stiffening in atherosclerosis (Figure E).

Altogether, our data indicates that TRPV4 contributes to matrix stiffening in atherosclerosis. Macrophages may not be the only source of cellular stiffness in atherosclerosis, as other cell types, including endothelial cells and pericytes, may differentiate into myofibroblasts during fibrosis development in atherogenesis. Further studies are needed to determine how TRPV4 regulates fibrosis, lesion development, and matrix stiffening in atherosclerosis and to explore the potential of therapeutically targeting TRPV4 to ameliorate atherosclerosis.

## Materials and Methods

### Mice and atherosclerosis model

Dr. Makato Suzuki (Jichi Medical University, Tochigi, Japan) initially generated TRPV4 knockout (KO) mice on a C57BL/6 background, later obtained by Dr. David X. Zhang (Medical College of Wisconsin, Milwaukee, WI). ApoE KO mice were sourced from the Jackson Laboratory, and ApoE:TRPV4 double knockout (DKO) mice on a C57BL/6 background were developed by the Rahaman lab. All mice were housed in a temperature- and humidity-controlled, germ-free environment with ad libitum food and water. All experimental procedures adhered to Institutional Animal Care and Use Committee guidelines and were approved by the University of Maryland’s review committee. For the atherosclerosis model, ApoE KO and ApoE:TRPV4 DKO mice were fed a high-fat diet (HFD; 21% fat, 0.15% cholesterol; Harlan Teklad) or a control chow diet for 12 weeks. Mice selected for thioglycolate-induced peritoneal macrophage collection were approximately 6-10 weeks old.

### Immunofluorescence staining

To detect TRPV4, CD68, and α-smooth muscle actin (α-SMA) proteins in human aortic root tissues, we obtained 5 μm sections from aortic root biopsies of atherosclerosis patients and healthy controls (OriGene; Rockville, MD). Double immunofluorescence staining was carried out to identify α-SMA+ myofibroblasts and CD68+ macrophages expressing TRPV4 (TRPV4+). Immunofluorescence intensity was quantified as integrated density. Tissue sections were incubated overnight at 4°C with primary IgGs—anti-CD68 (Bio-Rad), anti-TRPV4 (Alomone, Israel), and anti-SMA (Sigma-Aldrich)—at a 1:100 dilution to label TRPV4+ macrophages and myofibroblasts. Secondary staining was performed with Alexa Fluor 488- and 594-conjugated IgG (1:200), and DAPI was used to stain cell nuclei.

### Atomic force microscopy

We used the JPK Nano Wizard 4 atomic force microscope (AFM) (Bruker Nano GmbH, Berlin, Germany) to assess stiffness in aortic root tissues (human and mouse) and macrophages (5). For tissue samples, 7-10 μm-thick aortic root sections were mounted on glass slides and submerged in PBS, with individual force spectroscopy curves (F-D curves) recorded in contact mode. Measurements were conducted with CP-qp-CONT-BSG-B-5 colloidal probes (sQube), featuring a 30 kHz resonance frequency in air, a spring constant of 0.1 N/m, gold coating on the detector side, and a 10 μm borosilicate glass sphere. Parameters included a 1 nN setpoint, 10 μm/s extension speed, and a 10 μm Z length to obtain consistent F-D curves, with 50 curves recorded per sample across multiple regions. Young’s modulus (E) was calculated using the Hertz model for spherical probes, and data visualization, including histograms and plots, was performed using GraphPad Prism.

For macrophage stiffness measurements under live conditions, the AFM was operated in Quantitative Imaging (QI™) mode, allowing simultaneous nanomechanical characterization and imaging. Macrophages were seeded on poly-D-lysine-coated glass-bottom dishes (FD35PDL, WPI) and treated with or without LPS (100 ng/mL) for 24 hours. Measurements were conducted in PBS at 37°C with 5% CO_2_, and QI maps of 128 × 128 pixels were obtained. qpBioAC-CI-CB2 cantilevers (Nanosensors) with a 50 kHz resonance frequency in air, a 0.1 N/m spring constant, partial gold coating on the detector side, and a 30 nm-radius quartz probe were used. A total of 10 cells per condition were imaged, with 3-4 areas scanned per cell at an imaging setpoint of 0.5 nN.

## Data availability

All data generated or analyzed during this study are included in this article.

